# Small-scale field assessment of efficacy of the autodissemination approach against Aedes sp. in an urban area

**DOI:** 10.1101/2020.01.13.904094

**Authors:** Ahmad Mohiddin Mohd Ngesom, Hidayatulfathi Othman, Rawaida Bahauddin, Nazni Wasi Ahmad, Lee Han Lim, Asmalia Md Lasim, Yanfeng Liang, David Greenhalgh, Jasmine Chia Siew Min, Mazrura Sahani, Rozita Hod, Topek Omar

**Affiliations:** School of Applied Health Sciences, Faculty of Health Sciences, Universiti Kebangsaan Malaysia, Jalan Raja Muda A. Aziz, 50300 Kuala Lumpur, Malaysia; Medical Entomology Unit, Infectious Disease Research Centre, Institute for Medical Research, Jalan Pahang, 50588 Kuala Lumpur, Malaysia; Department of Community Health, Faculty of Medicine, Universiti Kebangsaan Malaysia, Jalan Yaacob Latif, Bandar Tun Razak, Cheras, 56000 Kuala Lumpur; Faculty of Science and Technology, Universiti Kebangsaan Malaysia, Bangi, 43600 Selangor, Malaysia; Department of Mathematics and Statistics, University of Strathclyde, 26 Richmond Street, Glasgow, G1 1XH United Kingdom; Department of Biomedical Sciences, Faculty of Medicine and Health Sciences, Universiti Putra Malaysia, 43400 UPM Serdang, Selangor Darul Ehsan, Malaysia; Disease Control Division, Ministry of Health Malaysia. Level 3, Block E10, Complex E, Federal Government Administrative Centre, 62590 Federal Territory Putrajaya, Malaysia

## Abstract

This is the first study to evaluate the efficacy of an autodissemination approach, as suggested by WHO. Therefore, the efficacy of an autodissemination approach in small-scale field trials against wild *Aedes* sp. population was evaluated in an urbanized setting, Malaysia. Lethal ovitraps enhanced with pyriproxyfen were used to control *Aedes* sp. populations at treatment sites, with the autodissemination activity was assessed using the WHO larval bioassays. Lethal ovitraps enhanced with pyriproxyfen effectively reduced of *Aedes* sp. population. All autodissemination stations were shown to be visited by *Aedes* sp. mosquitoes with 100% complete inhibition against eggs and larvae development. In the larvae bioassay, pupae mortality ranged from 14 to 40%. Statistically, a significant reduction of *Aedes* sp. population in the treatment sites compared to the untreated areas. The study proved for the autodissemination of pyriproxyfen to breeding habitats by wild *Aedes* sp. This technique is highly potentially for vector control activities. Future evaluation should focus on large-scale field trials.

**Author Summary:** Since 2012, Dataran Automobil, Seksyen 15, Shah Alam, was declared as one of the dengue hotspot areas. Major vector control activities were conducted by government, NGOs, social communities, and local authorities, but the number still rising. We conducted a new invention of autodissemination concepts in this area by an entomological study on mosquito populations reduction and dispersal abilities of the technique. We found that the technique has proven to control mosquito populations, but the other factors such as epidemiology link still unclear and need further clarification. Our finding highlighted the effectiveness of autodissemination strategies that can be considered as one of the alternative tools in vector control programme.

## Introduction

Dengue fever is a mosquito-borne disease, widely spread all over the world. The primary vector of dengue transmission is *Aedes aegypti* mosquitoes, while *Ae. albopictus* mosquitoes are secondary vectors. Both species also transmit yellow fever, zika, and chikungunya infections. Over the past 50 years, the spread of dengue infection worsened with a 30 times increase in global incidence cases. The World Health Organization (WHO) estimated around 50 – 100 million symptomatic dengue infections resulting from about 10,000 deaths annually. This puts half of the world’s population at risk in contracting dengue infection [1]. Dengue infection is considered as a global issue, with the Asian-Pacific region contributing to 75% of the current dengue cases [2]. The dramatic increase in dengue cases is most probably due to the growth in global human population as well as the unplanned urbanization of cities around the world. These lead to a rise in an ideal environment for dengue vectors [3].

Currently, there are no commercial vaccines and specific treatments for dengue infection. Thus, transmission of dengue fever is mainly controlled by prevention through vector control programs [4, 5]. *Aedes aegypti* and *Ae. albopictus* mosquitoes feed on human blood, typically active during the day. The immature larvae of these species can be found in a such as tires, artificial containers, can, bamboo stumps, and leaves. Among the control measures implemented against these species are through biological strategies (larvivorous fish and copepods), social mobilization, and chemicals [6]. Chemicals used in larvicide and adulticide target different life stages of mosquitoes. The use of larvicide was found to be more appropriate in controlling mosquito larvae or pupae. Larvicide targets the aquatic phase of the vectors, making it an advantage in controlling all the population of mosquitoes. However, the need for frequent application of larvicides at breeding sites has proven to be a limitation under operational conditions. Insecticides and labor required for large-scale vector control programs contribute to the significant cost required, which indirectly decreases the impact of the intervention as the frequency of such activities is low [7–11]. Hence, there remains an urgent need for new tools and strategies to reduce dengue transmission in a wide range of settings.

Pyriproxyfen (PPF) is a juvenile hormone analogue (JHA) that is effectively used in vector control programs, especially against mosquitoes. It inhibits the physiology of reproduction, morphogenesis, and embryogenesis of the insects [12]. The highly potent PPF larvicide inhibits mosquito emergence at a low dose, with no harm to human and non-targeted organisms. No resistance against pyriproxyfen has been reported in any mosquitoes. Besides, the larvicide activity of PPF can persist up to six months in different environments. This will affect the egg’s development, reduce the fecundity and fertilities of the eggs, and subsequently sterile the adult mosquitoes. The novel strategy of using mosquitoes to deliver pyriproxyfen to other breeding sites has become more interesting. It was first devised by Itoh et al. [13] and was then evaluated and improvised by other researchers on the coverage of the dissemination, doses used, the attractiveness of the formulation, and the efficacy of the autodissemination under semi and field evaluations [14–17]. Therefore, it is vital to identify the abilities of autodissemination of pyriproxyfen in Malaysia, which may ultimately affect vector control programs against dengue, zika, and chikungunya.

Here, we integrated data obtained from Shah Alam, Selangor, Malaysia, to assess the potential of pyriproxyfen autodissemination of controlling *Aedes* sp. in endemic dengue hotspots through monitoring of eggs, larvae and adult emergence in Shah Alam, Selangor. This study reports on the first experiments undertaken in a small-scale field condition, and we expect that result from this study will provide a better picture of autodissemination characteristic related to other parameters and factors.

## Methods

### Study sites

The study area (19°57’05” S, 43°76’88”) is an urban area in Shah Alam, Selangor. Dataran Automobil is located in Seksyen 15, Shah Alam, approximately 10-minutes by car from the Shah Alam Town Centre six kilometers away (Fig 1). This area consists of eighteen blocks of five-story shop lot, with each block separated by car parks and small roads. A small river flows down the side of the buildings with some natural and artificial vegetation shrubbery within the area. Due to its strategic location, Dataran Automobil is mostly occupied by foreign workers. This area was declared as a hotspot, with continuous dengue cases reported since 2012. There are multiple car parks, 24 shop lots, recreational parks, and two dumping sites areas. It is estimated that about 20 to 40% of the residents are immigrants from Nepal, India, Bangladesh, Cambodia, Indonesia, and China.

**Fig 1.**
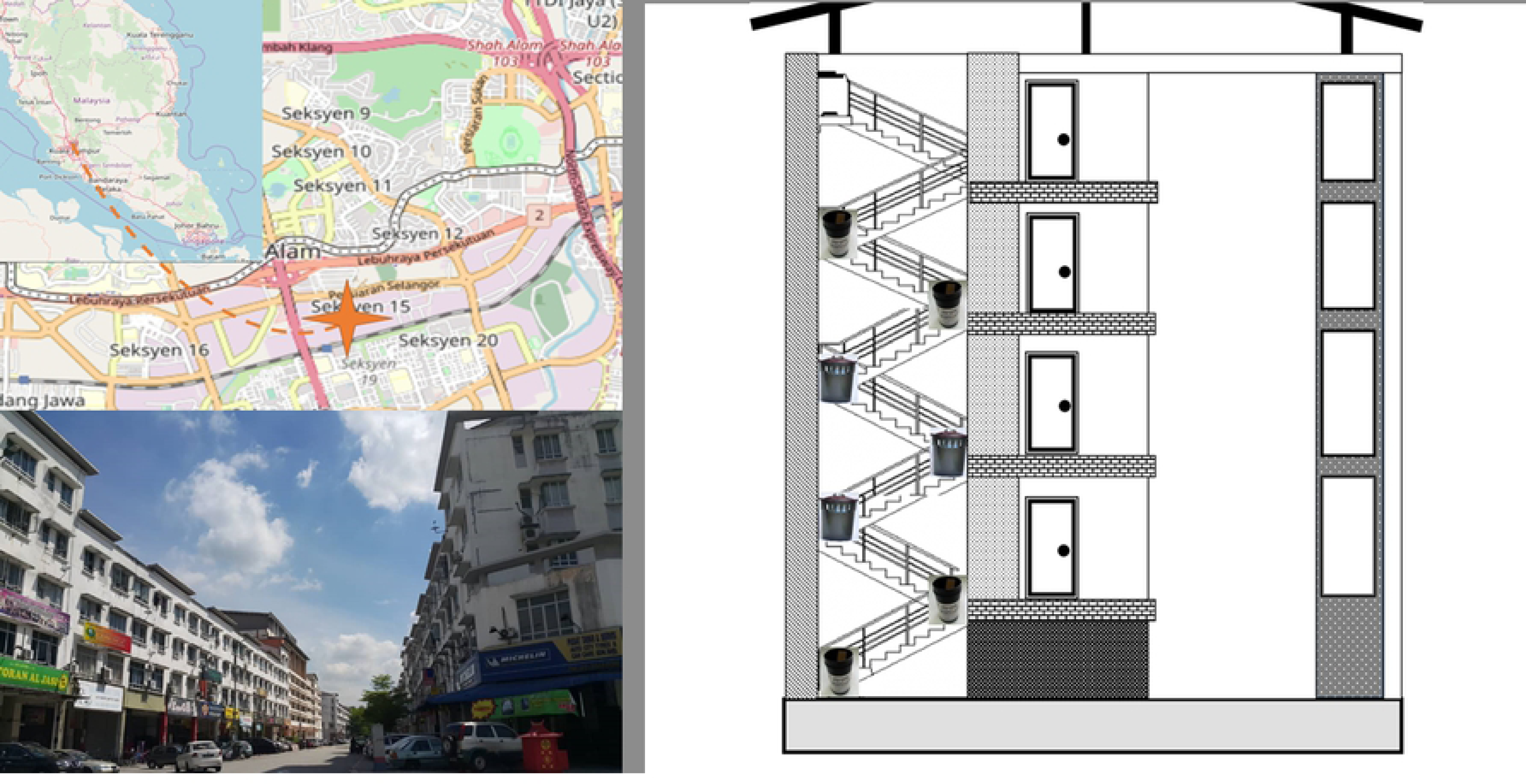
The location of study area, MHS, and ovitrap placement in Dataran Automobil, Seksyen 15, Shah Alam. The maps were obtained from U.S Geological; Survey, National Geospatial Program (https://glovis.usgs.gov/app?fullscreen=0; browsed on 4 January 2020) and the building photo was from courtesy of Ahmad Mohiddin.

### Mosquito Home Autodissemination

The Mosquito Home Autodissemination (hereafter referred as MHS) trap used for this research is described elsewhere [18]. Briefly, the device is a black polyethylene flowerpot with ten openings at the top of the device. The autodissemination device measured about 19.7cm (height) × 14.6 cm (diameter) in size. The application dosage of pyriproxyfen treated water is 0.004% w/w supplied by One Team Network, Sdn Bhd. The solution was tested in a room assay by the Vector Control Research Unit (VCRU), Universiti Sains Malaysia, by using oviposition cups and showed 100% mortality against all larvae in the experiments. During the trials, oviposition sites were placed in control and treatment sites, and autodissemination devices were only deployed in treatment sites. Autodissemination devices were serviced fortnightly to ensure all functions were working, sufficient water solution treatment, and removal of any clogged devices.

### Study Design

The trial procedure was slightly modified from previous protocols [19–21]. The study was conducted for duration of 9 months at Dataran Automobil, Seksyen 15, Shah Alam. The trial consists of two months prior to PPF dissemination, six months of citywide PPF dissemination, and one-month post-PPF dissemination. Initially, conventional ovitraps were deployed for the trials. The ovitrap was observed at weekly intervals, and larvae were identified to species to determine the ratio of *Aedes* sp. The data will provide the size of *Aedes* sp. population within the study area. In the third month, the Mosquito Home System (MHS) is deployed to selected treatment areas for six months. Following the MHS deployment, conventional ovitraps were placed in between MHS traps at a distance interval range of 1 – 5 meters within access of the *Aedes* population. The larvae density and ratio of *Aedes* sp. were recorded continuously until the end of the trials.

### Ovitrap Surveillance

A total of 160 oviposition cups (12 × 4 treatment sites, 12 × 4 control sites) (Fig 1) were placed at the stairways of eight blocks, along with the second and fourth levels. Ovitrap technique with paddles was used to monitor the *Aedes* populations and changed weekly as required [22]. Each ovitrap cup was coded individually with labels to ensure it is placed at the same location in each sampling round. Any missing ovitrap or paddle was replaced with a new one. The ovitraps were taken to the laboratory separately, and water from the ovitrap will be transferred to an enamel pan. All the larvae were identified using the Entomological Charts for Teaching provided by the Unit of Medical Entomology, Institute Medical Research (IMR), Kuala Lumpur.

### Intervention

A total of 150 MHS devices were deployed in four blocks on all stairways. Three autodissemination devices were placed diagonally clustering around each trap on the third floor. Autodissemination devices were one to five meters from the nearest ovitrap. Autodissemination device deployment took place from January 2018 to June 2018, coinciding with the rainy season. All MHS devices were removed from the field at the end of month eight. MHS was checked fortnightly throughout the six months of the trials to refill the solution and replace any loss on MHSs.

### Activity of pyriproxyfen in the field

All field samples from ovitrap and autodissemination devices (MHS) were brought back to the laboratory and then filtered to remove organic debris and wild mosquito populations. From each sample, 200ml water was transferred into a bioassay cup to determine pyriproxyfen activity. Twenty laboratory-reared third instars larvae were exposed to the field samples following larval bioassay procedure as described above. Pupal mortality, abnormal morphology, or coloration was recorded to show pyriproxyfen activity in the field samples. In some specimens, the contaminations of pyriproxyfen increased the larval development time of typically 8-9 days up to two weeks and subsequently causing death in the pupae stages.

### Pyriproxyfen bioassays

#### Mosquito rearing

Larvae of *Ae. aegypti* were obtained from a colony established at the Laboratory of Institute of Medical Research. Mosquito was reared, as described by Mohiddin et al. [23]. Larvae were fed with Tetramin© fish food and adult mosquitoes were fed with 10% of sucrose solutions.

#### Larval bioassay

Larval bioassay was conducted to assess the impact of pyriproxyfen on the field according to the WHO guidelines [24]. From ovitrap containers, 200ml of water samples were used for the study. The water samples were then tested with three different points of time- before, ongoing and after the treatment. All larvae bioassays were performed using late third instar larvae of Ae. aegypti. For the negative control, three cups were set up using tap water and 20 larvae per bioassay. Control treatments were treated with 199ml tap water and 1ml of ethanol. The mortality was recorded every 48h until the emergence of adult, while the larvae were also provided with food daily. Experiments were conducted at temperature of 26±2°C and 60±20% RH and preferably a photoperiod of 12h light followed by 12h dark.

#### Meteorological parameter

Data were collected within the same period of larval collection, starting from November 2017 to June 2018. Climatic data were obtained from the Malaysian Meteorological Department, Petaling Jaya station. The weekly means for rainfall and mean temperature in Shah Alam, Selangor, is shown in Fig 2.

**Fig 2.**
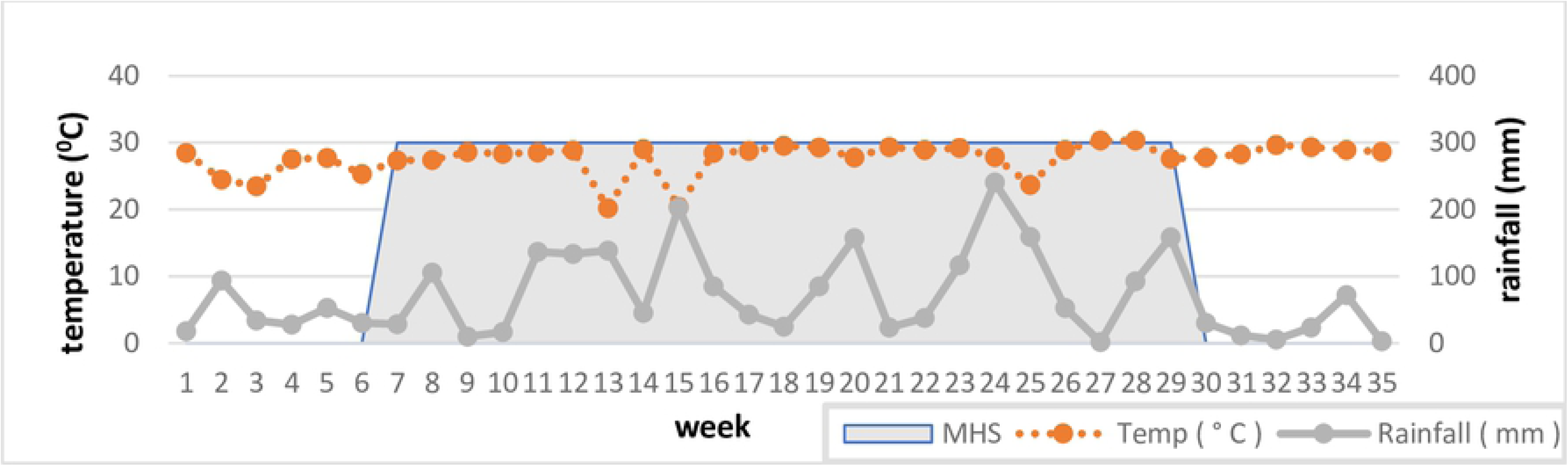
Weather variables of temperature (dashed line) and rainfall (solid line) obtained from the Dataran Automobil, Seksyen 15, Shah Alam.

### Data Analysis

The abundance of *Aedes* sp. mosquito populations was assessed using the positive ovitrap index, mean eggs per trap, and standard error were computed. The relationship between the meteorological variable and the number of *Aedes* sp. population was assessed with Pearson’s correlation coefficient. Pearson’s correlation coefficient was then used to express the correlation between the mean number of eggs collected from MHSs, percentage positive cup and percentage of pupae mortality. Pearson’s correlation coefficient test was also used to detect correlations between climatic factor and entomological parameter (number of eggs, larva) with a significant level set at 0.05. A generalized linear model (GLM) with Poisson distribution was carried out to compare the number of larvae population obtained from the ovitrap in treated and untreated areas. The response variables were the number of larvae populations of *Aedes* sp. mosquitoes. Site of the traps (1; if contained autodissemination and, 0; other), week and interaction of the week*site trapping act as predictors. As the time of sampling period was fixed, no offset was used. All statistical analysis in these studies was performed with Statistical Package for the Social Science (SPSS) version 23.

## Results

### Descriptive analysis of entomological data

A total of 113, 877 larvae mosquito were collected from Dataran Automobil Shah Alam in the 35 weeks (December 2017 to August 2018). *Ae. aegypti* represented 99.66% of all mosquitoes collected (*Ae. albopictus* was the only other mosquitoes observed). This species was encountered at both treatment and control sites. In treatment and control sites, 99.88% and 99.43% *Aedes* sp. larvae were identified as *Ae. aegypti*, respectively (Table 1). Furthermore, the mean number of larvae collected was slightly lower in treatment (1602.51±1077.56) compared to the control sites (1651.11±1031.99). On the other hand, the mean number of *Ae. albopictus* larvae were higher in treatment (2.27±0.84) than in the control sites (0.49±0.48).

**Table 1.**
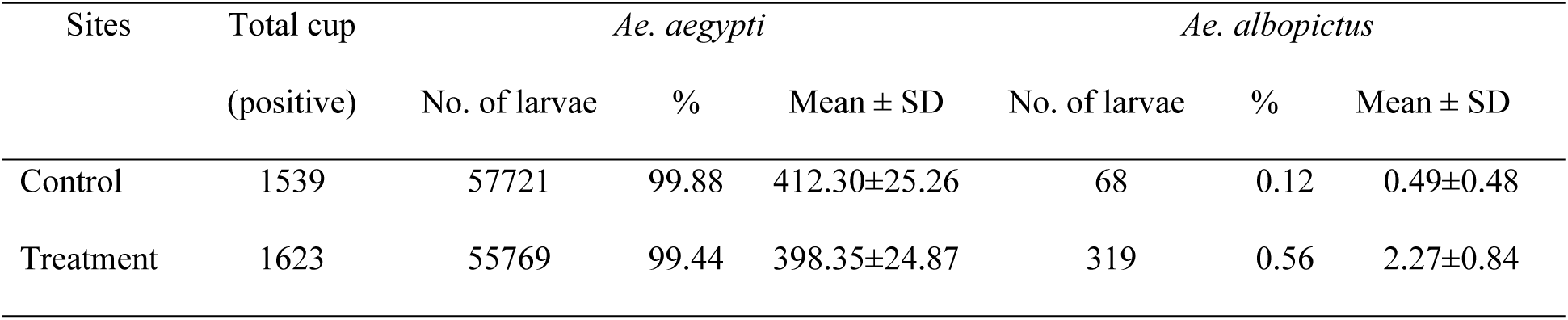
The overall number of *Ae. aegypti* and *Ae. albopictus* collected from control and treatment sites.

The number of single and mixed breeding among the *Aedes* sp. population in treatment and control sites were compared (Table 2). The mixed breeding of *Ae. aegypti* larvae were 1.96% and 6.8% obtained from control and treatment sites, respectively. The number of *Ae. albopictus* in mixed containers were 6.0 to 11.0 fold lower than *Ae. aegypti* species. Additionally, there was no significant difference (p=0.803) between mixed breeding sites to the mean number of larvae of these species in the treatment and control sites.

**Table 2.** Distribution of *Aedes* sp. based on single and mixed breeding sites from treatment and control areas.

| Sites | Single breeding |  |  |  |  |  | Mixed breeding |  |  |
| --- | --- | --- | --- | --- | --- | --- | --- | --- | --- |
|  | *Ae. aegypti* |  |  | *Ae. albopictus* |  |  |  |  |  |
| | No. of larvae | % | Mean ± SE | No. of larvae | % | Mean ± SD | No. of larvae | % | Mean ± SD |
| Control | 56853 | 98.38 | 411.98±25.62 | 0 | 0 | 0 | 936 | 1.62 | 468±22 |
| Treatment | 52276 | 93.21 | 408.41±26.87 | 0 | 0 | 0 | 3812 | 6.79 | 317.67±35.56 |
| Student *t*-test | | $P<0.005$ | | | nil | | | $P=0.803$ | |

### Influence of meteorology parameters/Description temporal

To understand the role of meteorological factors on *Aedes* sp. populations, the rainfall data and temperature were extracted from the study sites. Pearson’s correlation analysis showed a low association between the parameters selected. However, all parameters used in study areas did not show any significant associations between the meteorological and entomological parameters used (Table 3)

**Table 3.** Pearson correlation analysis between entomological parameters and meteorological data

| Meteorological<br>parameter | Entomological data |  |  |  |  |
| --- | --- | --- | --- | --- | --- |
|  | Rainfall | Ovitrap index | No. of larvae | No. of eggs | No. of eggs<br>(MHS) |
| Rainfall | 1 | -0.032 | -0.019 | -0.119 | 0.236 |
| Temperature | -0.431** | -0.126 | 0.276 | 0.128 | 0.101 |

\*\* denote significant correlations at the 0.05 and 0.01 significant levels.

### Impact of the autodissemination approach on the small-scale field

#### Autodissemination device (MHS)

The small-scale field evaluation provided information regarding the direct impact of pyriproxyfen on Aedes mosquito population. From the study, none of the larvae collected from autodissemination device solution emerged into adult. This result also showed 100% mortality against larva during the bioassay under laboratory conditions.

#### Ovitrap

Autodissemination of pyriproxyfen was detected in water collected from the ovitraps of the small-scale field trials. During the trials, the overall mortality of the pupa was 27.01±1.8%. Pupal mortality was 28.07±4.42% in week one post-treatment and was consistent during the trials and reached a peak in week 24 with 42.63±11.46%. A slight reduction in mortality was seen in week 12 (14.61±2.03). A different mean number of eggs collected from ovitraps during the study period ranges from 60.66 to 119.62 eggs per cup. The percentage mortalities of larvae showed a positive association with the mean number of eggs collected from MHS (r= 0.001) and with the positive cup of pyriproxyfen (r=0.238) (Fig 3). The higher number of eggs was related to the resting periods during the egg-laying, which ultimately enhances the possibilities to transfer more insecticide to other containers, thus increasing the number of mortalities against the larvae.

**Fig 3.**
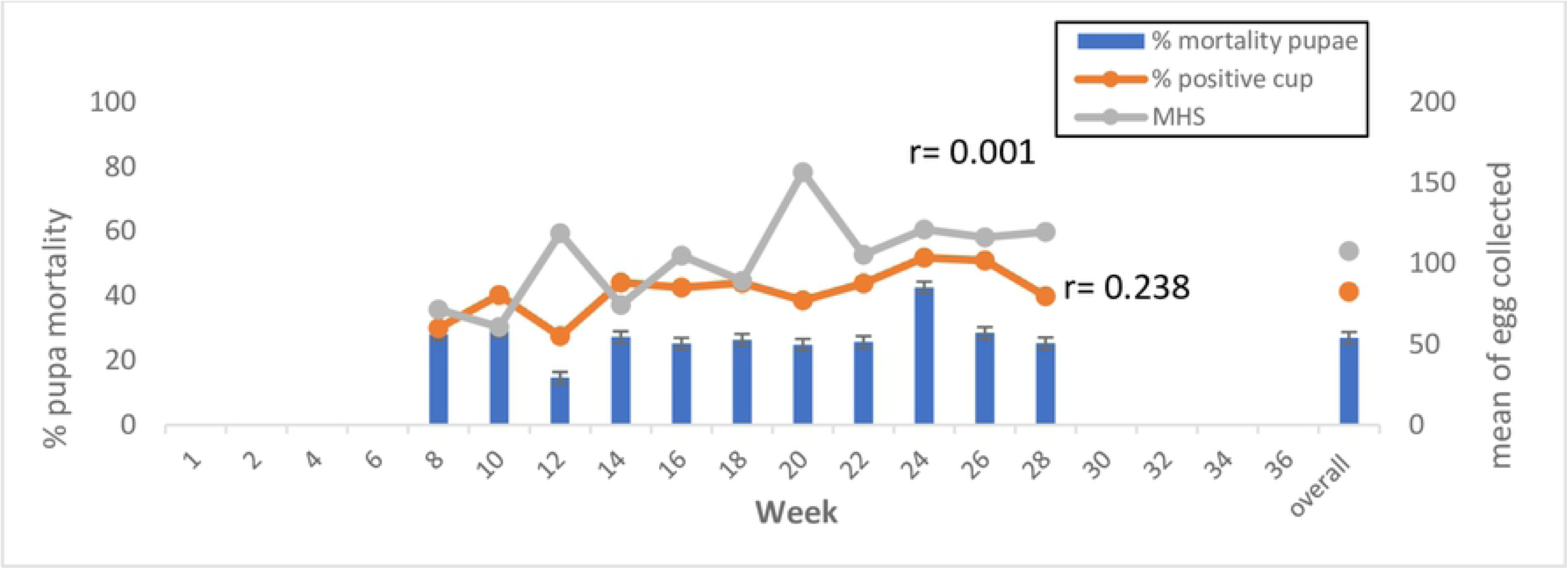
Pupa mortality percentage, % of positive cup and eggs collected by MHSs by week in treated sites

The Generalized Linear Model (GLM) with Poisson distribution used to analyze the data showed that the experiment’s sites are significantly predictors of the mean number of larvae (p<0.0001). The larvae populations were higher in the untreated sites compared to the treated sites during the implementation of autodissemination station. Unexpectedly, a significantly positive relationship between other predictors was observed (p<0.0001) (Table 4). Reduction in the number of larvae was observed in the early period of autodissemination deployment, constantly below untreated sites throughout the study period, with variations due to other factors (**Figure 4**).

**Fig 4.**
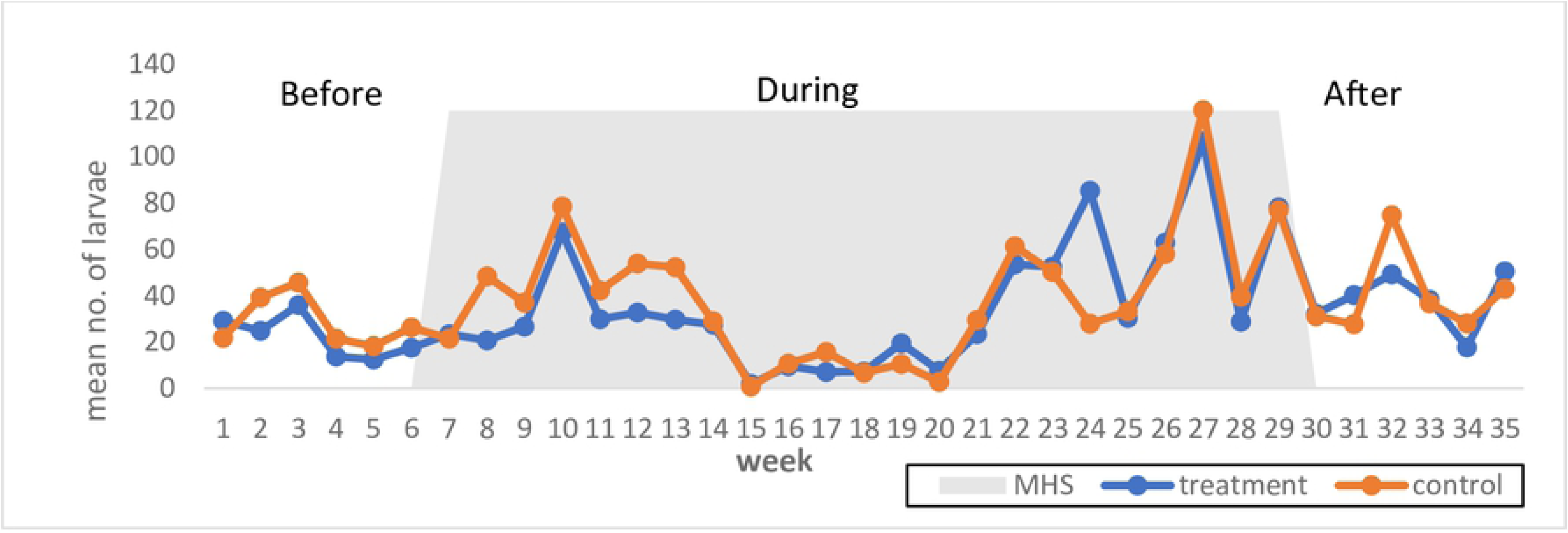
Larval population-based on autodissemination deployment in treated and untreated sites.

**Table 4.** *Aedes* sp. larvae abundance in treated and untreated sites using the Generalized Linear Model.

| Term | Estimate | SE | 95% CI |  | Pr(>\|z\|) |
| --- | --- | --- | --- | --- | --- |
| Intercept | 7.248 | 0.0171 | 7.215 | 7.282 | <0.0001 |
| Treatment | -0.812 | 0.0255 | -0.862 | 0.762 | <0.0001 |
| Week | 0.009 | 0.0009 | 0.008 | 0.011 | <0.0001 |
| Treatment*week | 0.042 | 0.0013 | 0.040 | 0.045 | <0.0001 |

Number of sampling:70, no. of week: 35, SE: standard error, Pr(|z|): statistic for z-value. ovitrap index as reference level.

## Discussion

This study is designed to assess the situation of a dengue epidemic area in Shah Alam, Malaysia, and investigate the effect of an autodissemination approach for vector control. The area was evaluated with entomological surveillance and climatic factors, while the autodissemination activity was assessed using the WHO larval bioassays. To the authors’ knowledge, no study has been conducted in Malaysia to determine the efficacy of an autodissemination approach under small-scale field trials. This approach has shown promising potential in reducing *Aedes* sp. population and thus improving vector control management during outbreaks, particularly in urban settings.

In the present study, using ovitrap technique, 118,391 eggs and 113,887 larvae were collected from the trial sites. *Ae. aegypti* was the predominant species found in almost all the treated and untreated sites. The ovitrap technique was found to assess the distribution and abundance of *Ae. aegypti* and *Ae. Albopictus* effectively, especially in areas with low frequencies of mosquito population. Ovitraps are widely used as large-scale monitoring concerning public health as they are inexpensive, easy to handle and highly sensitive compared to other techniques [25–26]. A large number of eggs and larvae found in this study could be due to the ideal environment surrounding Dataran Automobil in Shah Alam. Hence, the finding could be related to the larva-positive water tank, unmanaged rooftop structures, air-well design containing bottles, cups and plastic bags that provide suitable breeding conditions for *Aedes* sp. Other studies have also found that the poor design of rooftops and floors contributes towards a high number of breeding sites [27–28]. Recently, Garcia-Sanchez et al. [29] reported a water-tank containing detritus and planktonic organisms to be positive of mosquito larvae. The richness of bacterial community in the water-storage is directly/indirectly used as indicators for the presence of mosquitoes [30]. However, other factors should be taken into accounts, such as meteorological factors, vector control activities, and human activities. These factors taken together should be incorporated in dengue outbreak control programs.

Based on our analysis using data collected by mosquito population monitoring approaches, climatic factors may also contribute to the changes of mosquito population in the environment. Climatic factors are essential to the dynamics of mosquitoes in the epidemiological trend related to the transmission of diseases [31]. Temperature is usually associated with the effects on the viability of egg, larval development, longevity, and adult dispersal, whereas rainfalls affect the productivity of the mosquitoes and abundance of the species [32–33]. In San Luis, Brazil, the rainfall and three months’ lag of precipitation positively correlate with dengue cases reported [34]. Other findings with three different models show high significance between rainfall and *Aedes* populations [35]. Barrera et al. [36] found that adjusted rainfalls were also significantly and positively related to the number of adult mosquitoes collected from traps.

We also observed that temperature is an important variable in determining the *Aedes* sp. populations. Our findings show a positive association between temperature and the number of *Aedes* sp based on Pearson’s correlations analysis. Tsai et al. [37] found that critically low (as low as 13.8°C) and higher temperatures may decrease the mosquito populations, with regards to the mosquito’s fertility and longevity [38]. Other studies on diurnal temperature found a significant interaction between larval density and temperature settings. This could be due to the fluctuations, larval density and low-volume container parameters in predicting the effects. Although each factor individually influences the *Aedes* mosquito population, interaction between various factors will have a more significant impact on the final proportion of the mosquitoes [39–40]. The mathematical model of temperature transmission was used to integrate the data obtained from the laboratory tests. The results show that the highest peak of mosquito abundance with high transmission was suggested at 26°C to 29°C; however, transmission can also occur between 18°C to 34°C [41]. Similarly, like other findings [42], our results suggest that both temperature and rainfall are more likely to drive an additional change in mosquito populations, as well as a possible expansion in habitat for urban mosquitoes.

A high number of *Aedes* sp. populations in the environment may increase the probability of dengue transmission. A model was developed to estimate the abundance of mosquitoes in communities with their impact against dengue cases. The results demonstrated a definite risk of transmission resurgence due to the number of mosquito populations. However, the actual risk depends on the probabilities of uninfected people arriving at the right time of the year, where the mosquito population is at its peak with vectorial competence [43]. Most of the studies conducted relate the entomological indices with the number of cases and other related factors [44–46]. A study in Thailand investigated the relationship between sensitivity of entomological indices and immunological indicators in sites of treated and untreated areas [47]. Comprehensive research on mosquito abundance and dengue virus infection was also conducted in Peru. However, the results were not sensitive enough to detect any small changes in the transmission [48]. Ahmad et al. [49] agreed that high frequency of vector population control based on entomological indices results in a significant relationship between epidemiological and entomological parameters. It is crucial to expect high number of dengue cases in area with high density of *Aedes* sp. population and necessary preventive actions should be taken to avoid occurrence of dengue outbreaks.

Malaysia’s current vector control strategies still rely on conventional methods, which focus on thermal fogging, ultra-low volume spraying, larviciding, source reduction and enforcement activities [50]. Most of these strategies are not capable in solving the problem, which is primarily caused by larvae of *Aedes* sp. mosquitoes. Although thermal fogging eliminate adult mosquitoes, their progenies are still viable in breeding sites elsewhere. Unfortunately, most insecticide used for fogging has one-way effects [51]. In some areas, larviciding is the most appropriate method to control *Aedes* sp. population. However, workers often cannot find cryptic and hidden breeding sites during surveillance [52]. Based on these issues, new paradigms in vector control are required. An effective control measure against *Aedes* sp. mosquitoes is in need, which will be able to specifically target adult, larvae and cryptic breeding habitats of mosquitoes.

The autodissemination approach has become one of the most interesting methods to combine with other vector control strategies. The effectiveness of autodissemination is highly dependent on the *Aedes* sp. working as a form of transportation to disseminate insecticide to other breeding sites, in line with their “skip-oviposition” behavior [53–54]. In our study, all autodissemination stations were shown to be visited by *Aedes* sp. mosquitoes with 100% complete inhibition against eggs and larva development. Thus, MHS could be exploited to provide an index similar to other techniques (such an ovitrap) as it also assesses the percentage of positive breeding sites. However, results obtained from MHS containers are unable to calculate any mortality rates among the larvae as the eggs or first instar stage dies due to pyriproxyfen effects. Due to the new fortnightly setup of MHS formulation, we did not proceed with further evaluation against the mortality or any residual effect of the solutions. A commercial formulation of MHS with 0.004% active ingredient is lower than the concentrations used in other studies against dengue and malaria vectors.

The egg hatching rate is dose-dependent against insect growth regulators such as pyriproxyfen, diflubenzuron, and azadirachtin. Pyriproxyfen is considered as an ovicidal agent, especially on freshly laid eggs, which are more vulnerable and more sensitive than diapause eggs [55]. Suman et al. [56] found that higher concentration of pyriproxyfen exposure will significantly kill newly deposited and fully embryonated eggs. The hatching rate under non-diapausing conditions between newly deposited and fully embryonated eggs ranged from 1.0 to 1.8% against 1ppm pyriproxyfen. The efficacy of autodissemination traps does not rely on the hatching rate of larvae, considering the mortality rate as well as the presence of larvicides or insecticides. This finding was consistent with our result, whereby 100% larvae were killed and no emergence into adult was observed when we used 40ppm pyriproxyfen. Pyriproxyfen induced morphological abnormalities in *Aedes* sp. non-diapause embryos, leading to ovicidal effects. However, the mechanism of pyriproxyfen on egg diapauses in mosquitoes is still unclear [56–57].

For a system to have an autodissemination concept, it needs to have the ability to transfer the chemicals from the station to other breeding sites [17]. During our study, we were able to demonstrate the autodissemination concept of wild *Aedes* sp. mosquitoes, where pyriproxyfen solution from MHS station was disseminated to other breeding sites (ovitraps). Surprisingly, we found promising results as pupal mortality ranged from 14% to 40%, perhaps due to the placement of MHS autodissemination station in the study areas as well as the structure of the buildings. Most of the autodissemination stations and ovitraps were placed about two to five meters from each other. The autodissemination placement increased the chances of exposed mosquitoes to transfer a sufficient amount of pyriproxyfen to other larval habitats before the formulation evaporates [57–58]. Therefore, our autodissemination stations were the first and primary oviposition sites for mosquitoes, most of which were coming from the entrance of the buildings.

Semi-field and field trials have shown that *Aedes* sp. can transfer pyriproxyfen from treated containers to the other larval oviposition sites with a high range of emergence inhibition rates on the mosquitoes. Several factors might explain the results of this study. Suman et al. [14] obtained 50.4% pupal mortality against *Ae. albopictus* with dissemination range up to 200m distance in residential areas. Similarly, Lloyd and colleagues [53], also reported autodissemination activities assessed 200m from the autodissemination vases and can be effectively used for five weeks against *Ae. albopictus* larvae. Thus, it is likely that most studies use a large amount of pyriproxyfen to increase the impact of autodissemination under semi and field conditions [13–17, 59–60]

Additionally, we found a positive relationship between the number of eggs collected in autodissemination station (MHS) with percentage of a positive cup (r=0.238) and percentage of pupa mortality (r=0.001). Based on the findings by Chism and Apperson [61], the number of pupal mortality may have an association with the residence time during the egg-laying activities. The longer the time it took for mosquitoes to lay their eggs, the higher the amount of pyriproxyfen carried and transferred to other containers, thus increasing pupal mortality rate. A good relationship was obtained from various field trials with the percentage of pupal mortality and proportion of sentinel contamination to positively correlation (r=0.6) between the parameters [14]. Our recent finding through mathematical modeling and simulation found out that a sufficient number of MHS placed in the study area could reduced dengue cases up to 91% [62]. This indicates that the autodissemination station and Mosquito Home AQ used in this study have good potential in attracting wild mosquitoes and subsequently disseminate pyriproxyfen to other containers under natural environments.

Based on our results, there is no clear indicator of the reduced number of *Aedes* sp. population during the treatment period. However, pupa mortality during the laboratory bioassays and negative correlations were observed against eggs and larva densities throughout the deployment of autodissemination devices. Thus, the fluctuated trends of ovitrap index, larval and egg populations were observed at both treated and untreated sites. We noted that our trial is located in the middle of the city with high range of mosquito populations. High frequencies of mosquitoes from adjacent and other areas may freely migrate to the treated areas. These mosquitoes may potentially replace the local mosquito population [63], thus laying their eggs inside the ovitrap as their first choice before they reach any autodissemination station.

The occurrence of various factors has limited our ability to investigate a broader aspect against the local mosquito population. Additionally, incoming mosquitoes may also play an important role in increasing the coverage of pyriproxyfen dissemination, which, unfortunately, may lead to failure of certain experiments [64]. Oviposition preference of mosquitoes brings a complex implication in understanding local populations following usage of a new formulation of pyriproxyfen with a potent attractant developed by the manufacturer. Some studies report that *Aedes* sp. mosquitoes express oviposition behavior for specific types of attractants, such as bamboo leaf infusions [65], octenol [66] and yeast-produced CO_2_ [67]. These types of substrates should be taken into account when autodissemination stations are used as they may enhance the success rate of insecticide transference. Furthermore, all aspects of oviposition substrates that act as breeding site cues should be further explored to improve the efficacy of the stations.

## Conclusion

Result of the present work is consistent with other studies conducted on the effectiveness of an autodissemination approach based on pyriproxyfen treatment, either in the semi or field trials. This suggests that intervention based on the autodissemination technique may affect (depending on the site of applications) in reducing *Aedes* sp. population and, subsequently, inhibit the transmission of dengue viruses. Thus, this is the first pilot study to investigate the potential usage of an autodissemination approach for dengue control programme in Malaysia. We found that wild mosquitoes can transfer pyriproxyfen to other larval breeding containers, thus reducing *Aedes* sp. population. The effectiveness of autodissemination approach was also affected by various factors such as type of treatment either direct or autodissemination, coverage of the treatment, type of pyriproxyfen, geographical and meteorological parameters. However, before this approach can potentially be used, further studies are needed to improve the autodissemination efficacy, especially on formulation development, including mark-release-recaptured (MRR) trials to understand the dispersal and migration pattern of the mosquitoes prior to population reduction under field conditions.

## Acknowledgments

The authors are grateful to One Team Networks Sdn Bhd for providing necessary equipment and assistance during the field experiments. We would also like to thank Farihan from Universiti Kebangsaan Malaysia and Khadijah from Pejabat Kesihatan Daerah Petaling for providing mosquitoes for laboratory experiments. This research was partially supported under the Dengue Tech Challenge 2016 Grant, under the Newton-Ungku Omar Fund Partnership (AIM/PlaTCOM/HIP2/CCGF/2016/088).

## Authors contributions

NWA, HFO, LHL and AMMN conceived the idea, developed the design and protocol for this research. AMMN and RB performed data collection and experiments. AMMN, YL, DG, AML and TO analysed the data. HFO, MS and RH supervised the study. AMMN draft the original manuscript. HFO, NWA, JCSM, YL, DG review and editing manuscript to final form. All authors read and approved the final version of the manuscript.

